# Association of intrauterine alcohol exposure and offspring depression: A negative control analysis of maternal and partner consumption

**DOI:** 10.1101/307462

**Authors:** Kayleigh E Easey, Nicholas J Timpson, Marcus R Munafò

## Abstract

**Background:** Previous research has suggested that intrauterine alcohol exposure is associated with a variety of adverse outcomes in offspring. However, few studies have investigated its association with offspring internalising disorders in late adolescence.

**Methods:** Using data from the Avon Longitudinal Study of Parents and Children (ALSPAC), we investigated the associations of maternal drinking in pregnancy with offspring depression at age 18. We also examined partner drinking as a negative control for intrauterine exposure for comparison.

**Results:** Offspring of mothers that consumed any alcohol at 18 weeks gestation were at increased risk of having a diagnosis of depression (OR 1.15, 95% CI 1.00 to 1.32), but there was no clear evidence of association between partners alcohol consumption during pregnancy and increased risk of offspring depression (OR 0.90, 95% CI 0.78 to 1.04).

**Conclusions:** Maternal drinking in pregnancy was associated with increased risk of offspring depression at age 18. Residual confounding may explain this association, but the negative control comparison of paternal drinking provides some evidence that it may be causal, and this warrants further investigation.

## Introduction

Alcohol consumption during pregnancy is common, with up to 80% of expectant mothers in westernised countries reporting consuming alcohol when pregnant (O’Keeffe et al., 2015). This high percentage of women reporting alcohol use may be in part due to previous guidelines, which suggested that low levels of consumption are safe for the developing foetus. Until recently in the UK, for example, guidelines advised pregnant women to abstain from alcohol in the first three months of pregnancy; however, these guidelines also state that there is no evidence that a low level of alcohol use of 1-2 units (2 units being a 175ml glass of wine), no more than once or twice a week is linked to harm to the unborn child (NICE, 2008). Guidelines for alcohol use during pregnancy have only recently been updated to advise that women should abstain from alcohol consumption during their entire pregnancy (Health, 2016). This change is due in part to growing evidence that maternal alcohol consumption in pregnancy is associated with several negative offspring health outcomes.

It is well established that heavy alcohol use in pregnancy can cause foetal alcohol syndrome (Mukherjee, Hollins, & Turk, 2006), resulting in physical and cognitive impairments (Coles, Platzman, Lynch, & Freides, 2002; Gibbard, Wass, & Clarke, 2003; Guerri, Bazinet, & Riley, 2009). However, even at levels of alcohol consumption below that required for foetal alcohol syndrome, exposure to alcohol during gestation has been shown to be associated with detrimental outcomes in the offspring, such as being small for gestational age (Mamluk et al., 2017), and birth complications such as pre-eclampsia and placental abruption (Salihu et al., 2011), as well as behavioural outcomes such as increased risk of externalising disorders (Sayal et al., 2014) and internalising disorders (Sood et al., 2001; Walthall, O’Connor, & Paley, 2008). However, much research in this area has been conducted on offspring at an early age, with less research in older age groups to establish whether these associations persist into adulthood. One of the few studies to have used an older offspring age group suggested that the detrimental outcomes shown for gestational exposure to alcohol are likely to be permanent as they were still evident at age 22 (Day, Helsel, Sonon, & Goldschmidt, 2013), although replication of this finding is required. Low levels of intrauterine alcohol exposure have also been shown to be protective against offspring internalising and externalising problems in some studies, suggesting that residual confounding may influence observed associations (Kelly et al., 2009; Robinson et al., 2010).

Frequency, pattern and timing have also been shown to be important when investigating maternal alcohol use in pregnancy, as opposed to just the presence or absence of consuming alcohol (O’Leary et al., 2010). Day and colleagues reported a dose-response association for alcohol use during pregnancy across all three trimesters with increased offspring mental health problems (Day et al., 2013). However, the evidence is mixed for specific associations during different trimesters. Niclasen and colleagues reported evidence that binge drinking at both 16 and 30 weeks gestation is associated with conduct disorder (Niclasen, Andersen, Strandberg-Larsen, & Teasdale, 2014). On the other hand, O’Leary did not find evidence of an association with internalising disorders when their analyses were restricted to the third trimester (O’Leary et al., 2009).

Observational studies such as these can identify associations. However, well described problems of residual confounding introducing bias, means there is conflicting evidence on the effects of intrauterine alcohol exposure, and causal inference is difficult. Mendelian randomization (MR) is a method used to support causal inference regarding exposures and outcomes, which can go some way to protecting against the limitations from confounding factors often seen in association studies (Davey Smith & Ebrahim, 2004). MR has shown that alcohol metabolising genes, which affect the speed of ethanol metabolism (with a fast metabolism hypothesised to protect the developing offspring in utero) are related to low offspring IQ scores, for mothers with moderate alcohol consumption in pregnancy (Lewis et al., 2012). However, genetic variants identified for alcohol use to date have small effect sizes and might suffer from weak instrument bias, therefore reducing power to detect a true effect.

Negative control analyses are an alternative method to assess whether associations are due to confounding, or likely to be causal. This is done by using exposures or outcomes with similar confounding structures but no plausible biological link (Gage, Munafò, & Davey Smith, 2016). If an association is also shown in the negative control analyses, it is more likely to be due to confounding and not the original exposure of interest (Davey Smith, 2008). Negative control analyses have been previously used to investigate the effects of smoking during pregnancy on offspring mental health (Taylor et al., 2017). There are currently few studies using negative control analyses to investigate parental alcohol use during pregnancy, and these studies have mainly focused on offspring externalising disorders (Eilertsen et al., 2017) or IQ (Alati et al., 2008). We therefore sought to investigate associations between both the frequency and pattern of maternal drinking in pregnancy at 18 and 32 weeks gestation and offspring depression, using data from a population based longitudinal study. We also investigated whether any associations may reflect a causal effect, using negative control analyses of partner drinking in pregnancy on offspring depression, as both these exposures are likely to be influenced by similar confounding.

## Methods

### Sample

The Avon Longitudinal Study of Parents and Children (ALSPAC) is an ongoing population-based study, which recruited pregnant women residing in Avon, UK with expected dates of delivery between 1st April 1991 to 31st December 1992. The core sample consisted of 14,541 pregnant women, of which 14,062 were live births and alive at 1 year of age. Participants have been regularly followed up through clinic visits and questionnaires. Detailed information about ALSPAC is available on the study website which includes a fully searchable data-dictionary of available data (http://www.bris.ac.uk/alspac/researchers/data-access/data-dictionary). For further details on the cohort profile, representativeness, and phases of recruitment, see (Boyd et al., 2013; Fraser et al., 2013). Ethics approval for the study was obtained from the ALSPAC Ethics and Law Committee and the Local Research Ethics Committees.

### Measures

*Exposures*. Alcohol consumption during pregnancy was measured by: 1) Frequency of drinking: mothers and partners were asked separately the frequency and amount of alcohol consumed (within the past 3 months) at 18 weeks gestation. Response categories were never, <1 glass per week, 1+ glass per week, 1-2 glasses a day, 3-9 glasses a day and ≥10 glasses a day. For the analyses, the last two categories were combined to 3+ glasses a day. 2) Pattern of drinking (binge drinking): mothers and partners were asked the number of days they had consumed at least 4 units of alcohol (with examples given to indicate the equivalence of 1 unit in various types of alcohol). Mothers were asked at both 18 weeks (how many times within the last 3 months) and 32 weeks (how many times within the last month) gestation; partners were asked at 18 weeks gestation. Response categories were 0, 1-2 days, 3-4 days, 5-10 days, >10 days and every day. For our analyses, the last two categories were combined to >10 days.

*Outcomes*. Depression in offspring was measured using the computerised version of the Clinical Interview Schedule-Revised (CIS-R). The CIS-R is a computerised interview that is used to derive a diagnosis of depression (ICD-10 criteria). Depression was coded as a binary measure of depression.

*Confounders and sensitivity analysis*. Potential confounding factors associated with alcohol consumption and offspring psychiatric disorder were included in the analysis. Mother’s socioeconomic position (professional/managerial or other) measured during pregnancy, income (divided into quintiles) measured at age 3 and 4 years, home ownership (mortgage/non-mortgage) measured at 8 weeks gestation, marital status (married or not) measured at 8 weeks gestation, sex, parity (first born, 2+ born), maternal tobacco (yes/no) and illicit drug use (yes/no) in months 1-3 of pregnancy, and maternal depression at 18 weeks gestation (scores >12 highly associated with a diagnosis of depression) measured by the Edinburgh Postnatal Depression Scale (EPDS) (Cox, Chapman, Murray, & Jones, 1996).

### Statistical analyses

We used logistic regression to investigate associations between maternal and partner alcohol frequency (18 weeks gestation), binge drinking (18 and 32 weeks gestation) and a diagnosis of depression (CIS-R) at 18 years of age. Comparisons were made between the never drank controls in each alcohol exposure and each alcohol frequency/pattern group.

The impact of confounders on these associations was explored by comparing unadjusted estimates, to those adjusted for socioeconomic variables (model 1) and those further adjusted for maternal behaviour during pregnancy variables (model 2), and for partner alcohol use (frequency or pattern, dependant on exposure) during pregnancy (18 weeks gestation only) (model 3). By increasing the number of items adjusted for the sample size decreased, as individuals with missing data are excluded from analysis. Therefore, as a sensitivity analysis, all analyses were conducted of the full sample and then repeated only on participants with complete data.

Multiple imputation by chained equation (MICE) in Stata (Royston & White, 2011) was also used to generate a maximum dataset comprising of 100 imputed datasets, each with 10 cycles. Generation of more than one imputation model allowed for the uncertainty in predicting missing data, by adding variability to the imputed values in each dataset, which are then averaged together. The variability in results between each dataset reflect the uncertainty associated with the missing values, and using Rubin’s rules standard errors are calculated which account for the variability in these results (Sterne et al., 2009). By averaging the distribution of the missing data from the observed data, valid assumptions can be made which account for variability. This method assumes any systematic differences between the missing and observed values can be explained by differences in observed data and are Missing at Random (Sterne et al., 2009). Multiple auxiliary variables available from the ALSPAC cohort were used to assist in the imputation. These included the predictive factors used in the main analysis (e.g., socioeconomic position), as well as other measures related to the outcomes (e.g., EPDS), and earlier offspring depressive measures such Mood and Feelings Questionnaire (MFQ) (Angold et al., 1995).

Analyses were conducted using Stata version 14.2.

## Results

Overall, 14% of mothers (1834 of 13,195) reported drinking at least one alcoholic drink per week in the first three months of pregnancy. At 18 weeks gestation 9% of mothers reported (1192 of 13,149) binge drinking on 1-2 days within the past month. For mothers who provided information on alcohol frequency or pattern of drinking, 4191 and 4169 offspring respectively, provided information for CIS-R diagnosis of depression at age 18. Mother and offspring characteristics for full sample analysis are presented in Table 1. All further presented results are for imputed analyses unless stated otherwise.

**Table 1:**
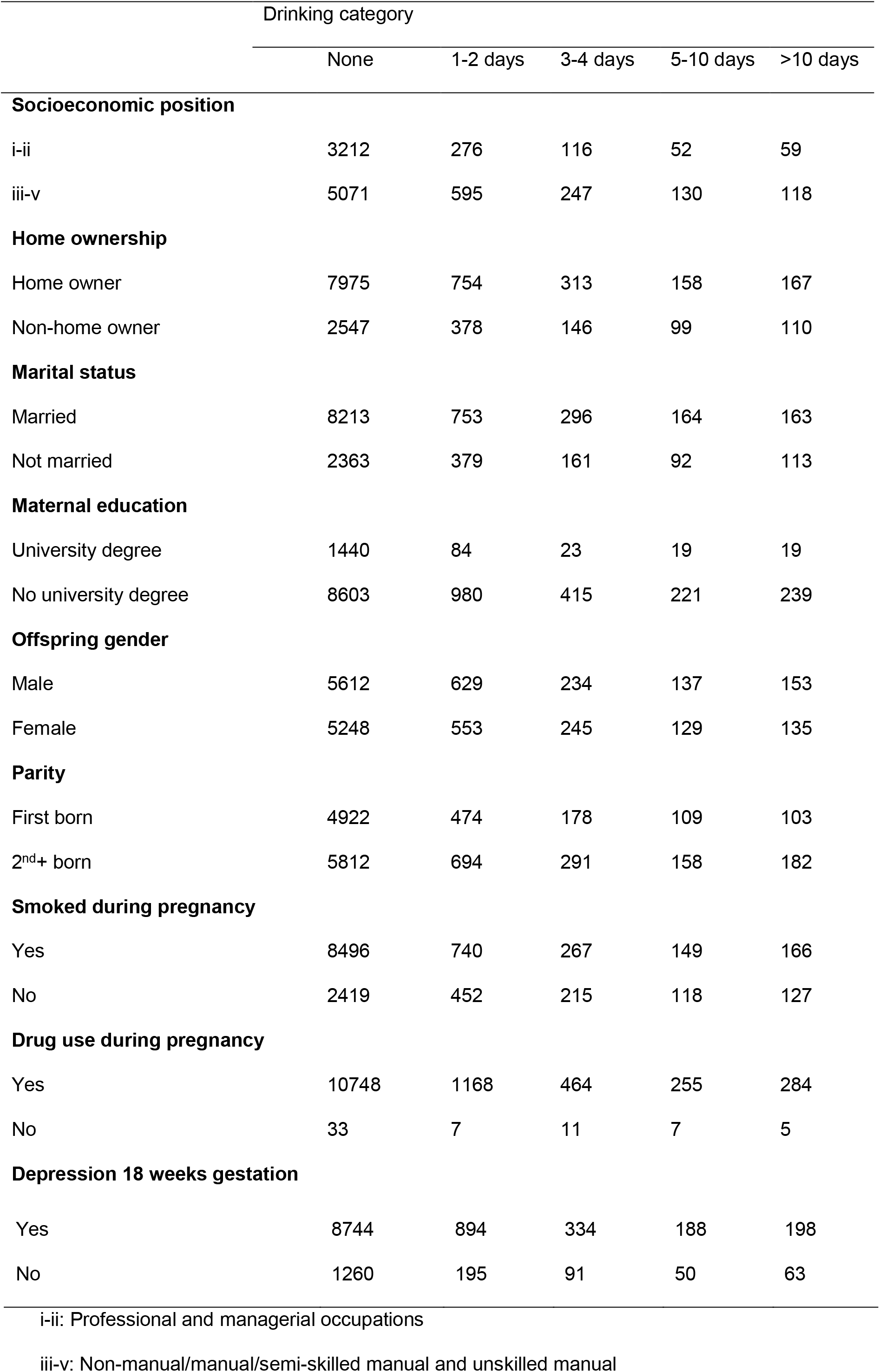
Mother and offspring socioeconomic factors, by pattern of drinking (number of times 4+ units of alcohol in past month at 18 weeks gestation)

|  | Drinking category |  |  |  |  |
| --- | --- | --- | --- | --- | --- |
|  | None | 1-2 days | 3-4 days | 5-10 days | >10 days |
| **Socioeconomic position** |  |  |  |  |  |
| i-ii | 3212 | 276 | 116 | 52 | 59 |
| iii-v | 5071 | 595 | 247 | 130 | 118 |
| **Home ownership** |  |  |  |  |  |
| Home owner | 7975 | 754 | 313 | 158 | 167 |
| Non-home owner | 2547 | 378 | 146 | 99 | 110 |
| **Marital status** |  |  |  |  |  |
| Married | 8213 | 753 | 296 | 164 | 163 |
| Not married | 2363 | 379 | 161 | 92 | 113 |
| **Maternal education** |  |  |  |  |  |
| University degree | 1440 | 84 | 23 | 19 | 19 |
| No university degree | 8603 | 980 | 415 | 221 | 239 |
| **Offspring gender** |  |  |  |  |  |
| Male | 5612 | 629 | 234 | 137 | 153 |
| Female | 5248 | 553 | 245 | 129 | 135 |
| **Parity** |  |  |  |  |  |
| First born | 4922 | 474 | 178 | 109 | 103 |
| 2<sup>nd</sup>+ born | 5812 | 694 | 291 | 158 | 182 |
| **Smoked during pregnancy** |  |  |  |  |  |
| Yes | 8496 | 740 | 267 | 149 | 166 |
| No | 2419 | 452 | 215 | 118 | 127 |
| **Drug use during pregnancy** |  |  |  |  |  |
| Yes | 10748 | 1168 | 464 | 255 | 284 |
| No | 33 | 7 | 11 | 7 | 5 |
| **Depression 18 weeks gestation** |  |  |  |  |  |
| Yes | 8744 | 894 | 334 | 188 | 198 |
| No | 1260 | 195 | 91 | 50 | 63 |

i-ii: Professional and managerial occupations
iii-v: Non-manual/manual/semi-skilled manual and unskilled manual

### Maternal alcohol consumption and offspring depression

Individuals whose mothers consumed any alcohol at 18 weeks gestation were at increased odds of having a diagnosis of depression at age 18 (unadjusted OR = 1.18, 95% CI 1.03 to 1.35). After full adjustment for socioeconomic and maternal behaviours these associations were attenuated only slightly (OR = 1.15, 95% CI 1.00 to 1.32, Table 2). Further adjustment for partner alcohol strengthened the association slightly (OR = 1.19, 95% CI 1.03 to 1.37).

**Table 2:**
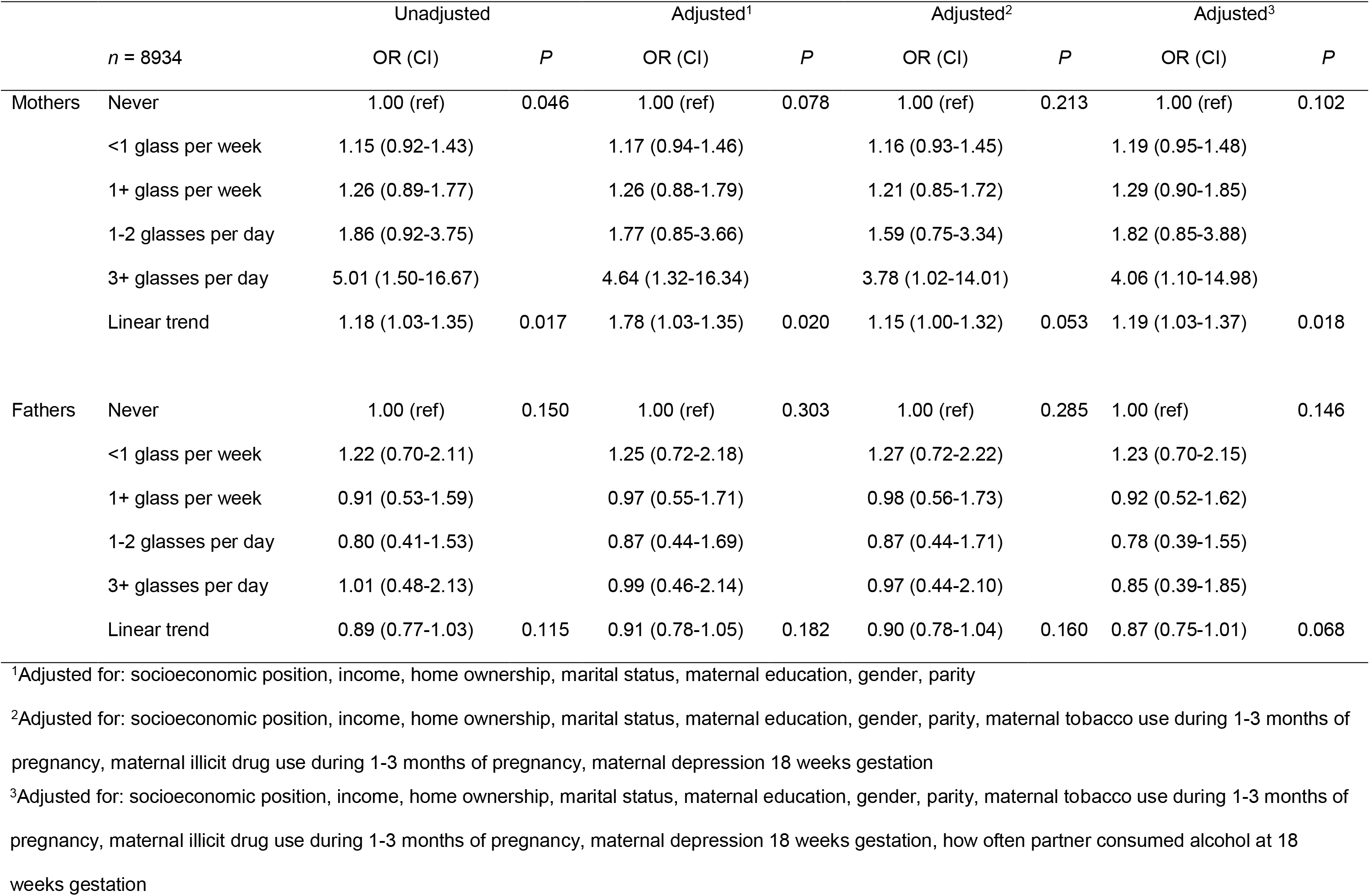
How often mothers and fathers consumed alcoholic drinks at 18 weeks gestation.

Consuming ≤4 alcoholic drinks at 18 weeks gestation was associated with a diagnosis of depression in offspring at age 18 (unadjusted OR = 1.14, 95% CI 1.01 to 1.29, Table 3). However, this association was attenuated after full adjustment for socioeconomic and maternal behaviours (OR = 1.06, 95% CI 0.93 to 1.21). Further adjustment for partner alcohol use strengthened the association slightly (OR = 1.08, 95% CI 0.94 to 1.23).

**Table 3:** Number of days mothers and fathers consumed 4+ units of alcohol in the past month (binges) at 18 weeks gestation.

|  |  | Unadjusted |  | Adjusted<sup>1</sup> |  | Adjusted<sup>2</sup> |  | Adjusted<sup>3</sup> |  |
| --- | --- | --- | --- | --- | --- | --- | --- | --- | --- |
|  | *n* = 8934 | OR (CI) | *P* | OR (CI) | *P* | OR (CI) | *P* | OR (CI) | *P* |
| Mothers | None | 1.00 (ref) | 0.171 | 1.00 (ref) | 0.465 | 1.00 (ref) | 0.653 | 1.00 (ref) | 0.579 |
|  | 1-2 days | 1.20 (0.83-1.73) |  | 1.15 (0.79-1.68) |  | 1.11 (0.76-1.62) |  | 1.13 (0.77-1.65) |  |
|  | 3-4 days | 1.77 (1.08-2.91) |  | 1.57 (0.94-2.60) |  | 1.47 (0.88-2.45) |  | 1.51 (0.90-2.53) |  |
|  | 5-10 days | 1.43 (0.65-3.14) |  | 1.29 (0.58-2.88) |  | 1.21 (0.54-2.74) |  | 1.27 (0.60-2.89) |  |
|  | >10 days | 1.23 (0.58-2.58) |  | 1.02 (0.47-2.18) |  | 0.92 (0.42-2.01) |  | 0.96 (0.44-2.11) |  |
|  | Linear trend | 1.14 (1.01-1.29) | 0.031 | 1.09 (0.96-1.23) | 0.169 | 1.06 (0.93-1.21) | 0.358 | 1.08 (0.94-1.23) | 0.274 |
| Fathers | None | 1.00 (ref) | 0.757 | 1.00 (ref) | 0.706 | 1.00 (ref) | 0.659 | 1.00 (ref) | 0.607 |
|  | 1-2 days | 0.91 (0.64-1.29) |  | 0.93 (0.65-1.33) |  | 0.93 (0.65-1.33) |  | 0.92 (0.64-1.32) |  |
|  | 3-4 days | 0.82 (0.59-1.15) |  | 0.84 (0.60-1.19) |  | 0.84 (0.59-1.19) |  | 0.83 (0.58-1.17) |  |
|  | 5-10 days | 0.96 (0.70-1.32) |  | 0.99 (0.71-1.37) |  | 0.97 (0.70-1.35) |  | 0.95 (0.68-1.33) |  |
|  | >10 days | 0.83 (0.57-1.23) |  | 0.81 (0.54-1.21) |  | 0.79 (0.53-1.19) |  | 0.77 (0.51-1.16) |  |
|  | Linear trend | 0.97 (0.89-1.06) | 0.504 | 0.97 (0.89-1.06) | 0.456 | 0.96 (0.88-1.05) | 0.383 | 0.96 (0.87-1.04) | 0.314 |

<sup>1</sup>Adjusted for: socioeconomic position, income, home ownership, marital status, maternal education, gender, parity
<sup>2</sup>Adjusted for: socioeconomic position, income, home ownership, marital status, maternal education, gender, parity, maternal tobacco use during 1-3 months of pregnancy, maternal illicit drug use during 1-3 months of pregnancy, maternal depression 18 weeks gestation
<sup>3</sup>Adjusted for: socioeconomic position, income, home ownership, marital status, maternal education, gender, parity, maternal tobacco use during 1-3 months of pregnancy, maternal illicit drug use during 1-3 months of pregnancy, maternal depression 18 weeks gestation, how often partner consumed alcohol at 18 weeks gestation

There was no clear evidence that consuming ≥4 alcoholic drinks at 32 weeks gestation was associated with offspring depression (Supplementary Tables 1 and 4).

### Partner alcohol consumption and offspring depression

Paternal alcohol use at 18 weeks gestation showed no clear evidence of association with offspring depression, for both frequency (unadjusted OR = 0.89, 95% CI 0.77 to 1.03, Table 2) and pattern of alcohol use (unadjusted OR = 0.97, 95% CI 0.89 to 1.06, Table 3). Further adjustment for mothers’ alcohol use slightly attenuated the strength of association, for both frequency (OR = 0.87, 95% CI 0.75 to 1.01, Table 2) and pattern of alcohol use (OR = 0.96, 95% CI 0.87 to 1.04, Table 3).

The findings from the full sample and complete case analyses did not differ substantially from the imputed analyses (see Supplementary Tables 2-11).

## Discussion

We investigated the associations between maternal alcohol consumption in pregnancy (frequency and pattern) and offspring depression in a population-based study.

Our results suggest that the amount of alcohol mothers consumed during pregnancy at 18 weeks gestation is associated with offspring depression at age 18, and indicate a dose-response relationship between maternal alcohol consumption in pregnancy and offspring risk of depression. Neither the amount of alcohol consumed, nor pattern of drinking by partners during pregnancy, were associated with offspring depression; indeed, the point estimates shown for the negative control analyses suggested associations in the opposite direction for maternal and partner drinking on offspring depression. The strength of associations shown for partner drinking and offspring risk of depression is almost as strong as maternal drinking, however, it is much less precisely estimated. This suggests that the associations shown for maternal alcohol consumption in pregnancy and offspring depression may be causal, and these associations are unlikely to be due to the result of shared confounding structures between maternal and paternal exposures. Such findings have implications for women trying to conceive or who may not be aware that they are already pregnant, when the foetus is most likely to be exposed to alcohol (Floyd, Decouflé, & Hungerford, 1999).

The pattern of maternal alcohol consumption in pregnancy at 18 weeks gestation indicated that increased episodes of maternal binge drinking were associated with increased risk of offspring depression. As binge drinking was defined by drinking four or more drinks on any one occasion whilst pregnant, these findings have implications for mothers who may drink heavily even only on one occasion whilst pregnant.

We did not observe any clear evidence of an association between binge drinking at 32 weeks gestation and offspring depression. This supports previous studies suggesting differences between trimesters for associations between maternal alcohol consumption and offspring internalising disorders, with risk of offspring mental health disorders not increased when analyses are restricted to third trimester alcohol exposures (O’Leary et al., 2009). Previous studies using the ALSPAC cohort have shown that the majority of women who binge drink during the earlier stages of pregnancy also report binge drinking during the later stages (Sayal et al., 2014), suggesting patterns of mothers alcohol consumption during pregnancy are relatively stable across trimesters. The difference in associations shown in the current study are therefore unlikely to be due to changes in mothers drinking patterns, but possibly in how alcohol may affect the developing foetus at different developmental stages.

Our use of a negative control comparison of paternal drinking in pregnancy provides some support that the observations we have observed may be causal. By using such a long follow up for the outcome measurement, the findings from the current study suggest that any effects shown within the offspring with intrauterine alcohol exposure are likely to persist throughout childhood and into adulthood.

Although the associations we observed are small, they may nevertheless be important at a population level, particularly as depression is a common mental health disorder affecting more than 300 million people globally (WHO, 2017). The population-attributable fraction (PAF) was calculated for alcohol exposure using UK prevalence of alcohol use during pregnancy from a recent meta-analysis (Popova, Lange, Probst, Gmel, & Rehm, 2017) for the comparison numerators and denominators. The PAF for the contribution of alcohol use in pregnancy on offspring depression was 0.054, suggesting if the associations we observed are causal and precisely estimated, we could potentially prevent up to 5% of depression cases by reducing alcohol consumption in pregnancy. Our findings therefore provide support for guidelines recommending complete abstinence from alcohol during pregnancy, or for women trying to conceive.

An advantage of the ALSPAC cohort, where recruitment occurred during 1990-1991, is that attitudes in the UK towards drinking in pregnancy were likely to have been different to current day with less stigma associated with drinking in pregnancy, meaning that mothers may have been more likely to truthfully report alcohol consumption. However, underreporting of alcohol use may still have occurred if mothers were not aware they were pregnant until later stages of pregnancy, therefore misrepresenting the true level of alcohol exposure as the alcohol exposures rely on valid self-report. If alcohol use was underreported, the findings we observed are likely to be more conservative and a larger association could have been shown if there was a more biologically valid way to assess maternal alcohol consumption.

There are limitations that should be considered when interpreting these results. Firstly, there is sample attrition from enrolment to the outcome measurement at age 18. Characteristics between responders and non-responders in the ALSPAC study could cause selection bias. However, as the complete case analyses and those using the imputed dataset do not differ substantially, selection bias is unlikely to have affected the reported associations. Previous studies investigating biases within the ALSPAC cohort have found the strength of associations to not be greatly affected by selection bias and sample attrition (Wolke et al., 2009). Secondly, the associations may also be due to a shared genetic risk for depression, which is expressed as different phenotypes in mothers and offspring. However, genetic data were not included in this analysis to be able to test this.

Our study highlights the potentially long-lasting detrimental effects of maternal alcohol consumption in pregnancy on offspring mental health. Although the associations we observed are small, they may nevertheless be important at a population level. The negative control comparison of paternal alcohol use during gestation provides some evidence that the associations found may be causal. However, further research is needed to determine with greater confidence whether these associations are indeed causal, and the result of intrauterine exposure. This may require the use of other methodological approaches, such as Mendelian randomization, and sibling comparisons with offspring discordant for maternal alcohol consumption in pregnancy.

#### Key Points

- High levels of maternal alcohol use during pregnancy has been associated with childhood mental health problems.
- There are contradictory findings on if light to moderate drinking in pregnancy is associated with offspring mental health problems.
- This study reports that light to moderate maternal alcohol use during pregnancy is associated with offspring depression at age 18.
- Our use of paternal alcohol use in pregnancy as a negative control, provides evidence that this association may be causal.
- This study offers support for total abstinence from alcohol during pregnancy.

## Acknowledgements and funding support

The UK Medical Research Council and the Wellcome Trust (Grant ref: 102215/2/13/2) and the University of Bristol provide core support for ALSPAC. We are extremely grateful to all the families who took part in this study, the midwives for their help in recruiting them, and the whole ALSPAC team, which includes interviewers, computer and laboratory technicians, clerical workers, research scientists, volunteers, managers, receptionists and nurses.

NJT is a Wellcome Trust Investigator (202802/Z/16/Z), a programme lead in the MRC Integrative Epidemiology Unit (MC_UU_12013/3) and works within the University of Bristol NIHR Biomedical Research Centre (BRC) and CRUK Integrative Cancer Epidemiology Programme (C18281/A19169).

KEE, NJT and MRM work in the Medical Research Council Integrative Epidemiology Unit at the University of Bristol which is supported by the Medical Research Council and the University of Bristol (this funds KEE’s PhD studentship).

KEE has full access to all the data in the study and takes responsibility for the integrity of the data and the accuracy of the data analysis.

## Ethical considerations

Ethical approval for the ALSPAC cohort study was obtained from the ALSPAC Ethics and Law Committee and the Local Research Ethics Committees.

## Corresponding author

Kayleigh E Easey, Tobacco and Alcohol Research Group, School of Experimental Psychology, MRC Integrative Epidemiology Unit, University of Bristol, BS8 1TU, UK.

